# Regulation of phage lambda packaging motor-DNA interactions: Nucleotide independent and dependent gripping and friction

**DOI:** 10.1101/2022.09.24.509349

**Authors:** Brandon Rawson, Mariam Ordyan, Qin Yang, Jean Sippy, Michael Feiss, Carlos E. Catalano, Douglas E. Smith

**Author notes:** Corresponding Authors: Douglas E. Smith, University of California San Diego, Department of Physics, 9500 Gilman Drive, La Jolla CA 92093; Carlos E. Catalano, University of Colorado Anchutz Medical Campus, Department of Pharmaceutical Sciences, 4200 E. Montlake Blvd, Aurora, CO, 80045.

## Abstract

Many dsDNA viruses utilize ATP-powered “terminase” motors to package their genomes into procapsid shells. Here we use a single-molecule DNA grip/slip assay with rapid solution exchange to probe effects of nucleotide binding/dissociation in phage lambda motors containing both the large (TerL) and small (TerS) terminase subunits. Both subunits are required for packaging in vivo, but for some viruses (e.g., phages T4, HK97) packaging can be measured in vitro with only the catalytic TerL subunit. TerS facilitates initiation of packaging in vivo, but it has remained unclear if it plays any role during translocation. Surprisingly we measure frequent DNA gripping and high motor-DNA friction even in the absence of nucleotide. Such behavior was not observed in phage T4 motors containing only TerL, for which motor-DNA interactions were measured to be much weaker and significant gripping and friction was only observed with nucleotide present. For the lambda TerL/TerS holoenzyme, binding of nucleotide (ATP analogs or ADP) further increases gripping and friction, indicating there are both nucleotide independent and dependent interactions. Our findings suggest that TerS plays an important role in motor processivity, and that ATP-independent DNA gripping explains pausing observed during lambda packaging. We propose TerS acts as a “sliding clamp” to limit back slipping when TerL loses grip. Additionally, we show that the lambda packaging complex has a “DNA end clamp” mechanism that prevents the viral genome from completely exiting the capsid once packaging has initiated.

## INTRODUCTION

Complex dsDNA viruses, including many bacterial viruses (phages) and human viruses such as herpesviruses, employ ATP-powered molecular motors to package their genomes into pre-assembled procapsid shells^1–6^. The motors exert large forces to pack DNA to high density, overcoming resistive forces arising from factors including electrostatic self-repulsion of DNA, DNA bending rigidity, hydration forces, and entropy loss^7–13^. These motors are also of interest as models for investigating general biomotor principles since they have homologies with many other cellular motors, including helicases, chromosome transporters, and protein translocases^14–16^.

DNA replication in most complex dsDNA viruses (tailed phages and herpesviruses) produces concatemers containing multiple genomes covalently linked head-to-tail from which individual genomes must be excised and packaged^1, 2, 5, 17^. Terminase enzymes perform both functions. The holoenzymes are hetero-oligomers composed of small TerS subunits that mediate specific recognition of viral DNA and large TerL subunits that possess the nuclease, ATPase and packaging (DNA translocation) activities (See **Figure 1**). While both subunits are essential for genome packaging *in vivo*, the two subunits do not strongly interact *in vitro*^1, 18–20^. Thus, many studies of viral motors have examined the large subunit in the absence of TerS. For instance, structural studies of the phage T4 motor composed only of TerL and the related phage ϕ29 gp16 packaging ATPase (which does not have a TerS subunit) indicate that motor units form a pentameric ring structure surrounding DNA^21, 22^. Lambda terminase, however, can be purified as a stable heterotrimer composed of TerL tightly associated with two TerS subunits (the protomer), which reversibly assemble into a pentameric ring-shaped holoenzyme complex [(TerL•TerS_2_)_5_] that possesses native catalytic activities ^23–26^. The arrangement of the TerS subunits in this complex form a decameric ring sufficiently large to encircle dsDNA, similar to ring structures that have been observed in other phage TerS complexes assembled in the absence of TerL^17, 18, 27^.

**Fig. 1.**
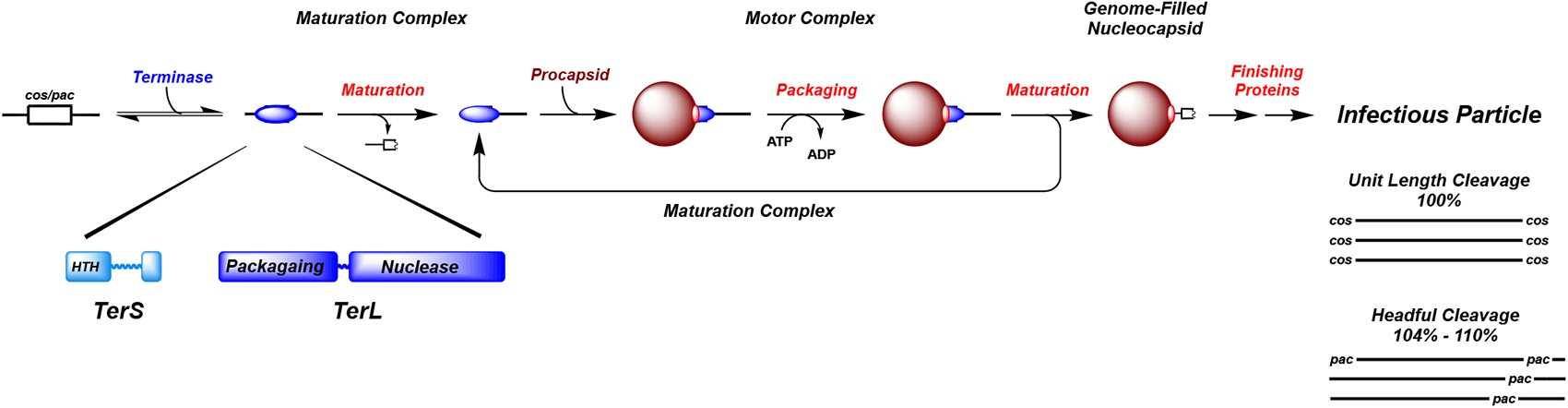
Genome Packaging in the Complex Double-Stranded DNA Viruses. Terminase enzymes are responsible for processive excision of an individual genome from a concatemeric packaging substrate (genome maturation) and for translocation of the duplex into a pre-formed procapsid shell (genome packaging). Two basic strategies for genome packaging, unit-length and headful, are summarized at bottom, right. Details are provided in the text.

Two distinct excision/packaging strategies have been identified in the dsDNA viruses (**Figure 1**)^1, 2, 4, 6^. Phages lambda, HK97 and T7, and the eukaryotic herpesviruses are examples of “unit length” genome packaging viruses. In these cases, packaging initiates at a specific site (“*cos”* in phage lambda), which represents the inter-genomic junction in the concatemer; specific assembly of terminase at *cos* is mediated by TerS. The nuclease activity of TerL cuts the duplex to “mature” the genome end and the terminase•DNA complex then binds to the portal of a procapsid shell. DNA is then translocated into the capsid, powered by the packaging ATPase in TerL, which continues until the next *cos* site is encountered at which point the motor stops and again cuts the duplex to release the nucleocapsid filled with a unit-length (100%) genome. The TerS subunit is responsible for specific recognition of *cos* during initiation and termination of packaging^25^. In contrast, phages P22, SPP1, and T4 are examples of “headful” packaging viruses. In these systems, packaging usually initiates at a “*pac*” sequence in the concatemer where the TerL subunit cuts the duplex, binds to a procapsid, and begins DNA packaging. Translocation continues until a head (capsid) full of DNA is packaged (104-110% genome length) and the DNA is then nonspecifically cleaved to terminate the packaging reaction (**Figure 1**). While TerS is required to initiate packaging *in vivo*, it is unclear what role it plays in termination. Moreover, in both strategies, the role that TerS plays in motor function during translocation remains unclear.

*In vitro* DNA packaging assays using defined components have been developed for phages phi29, lambda, T4, P23-45, and HK97^28–32^. Studies to date have found that TerS is required for lambda, modestly enhances packaging in HK97, but inhibits *in vitro* packaging in T4. Thus, at present, lambda is the most amenable system for us to study both TerL and TerS subunits in a holoenzyme complex. Importantly, all the bulk *in vitro* studies employ bulk DNase protection assays that measure overall efficiency of packaging but cannot distinguish between the effects on efficiencies or kinetics of packaging initiation, DNA translocation or termination/cleavage or retention of the packaged genome.

The development of methods using optical tweezers to measure the packaging of single DNA molecules into single phage capsids has facilitated more detailed studies of the translocation function of the phi29, T4, and lambda motors^8, 10, 33–35^. All three were found to generate remarkably high forces (>60 pN) and translocate DNA at rates as high as ∼200-2000 bp/s, depending on the virus. Studies with phi29 that resolved the incremental translocation steps of the motor and studies with T4 and lambda explored effects of site-directed residue changes in TerL have shed light on many aspects of the motor function and mechanisms^36–43^. However, much remains to be understood about the nature of motor-DNA interactions and the full range of factors that regulate them. The molecular interactions that allow the motor to grip the DNA are unclear, and the causes of features such as the stochastic pauses and slips that frequently occur during DNA translocation are not fully understood.

Studies and models suggest that ATP binding induces TerL to grip DNA tightly, while the release of the hydrolysis products (ADP/ P_i_) facilitates release of the grip^33, 41, 44^. This is supported by findings that an actively packaging motor can be stalled with the addition of poorly hydrolyzed ATP analogs (e.g., ATP-γS, (β,γ-imido)-ATP or (β-γ-methylene)-ATP, collectively referred to herein as ATP*), and that slipping during packaging increases when ATP concentration is lowered or when residue changes are made in the TerL ATP binding motifs^8, 33, 41, 45^. Ensemble biochemical assays with phage lambda found that DNA release from the stalled, partially filled capsids is very slow, suggesting that the motor usually retains a tight grip in the presence of nucleotide^46^.

To probe the dynamics of motor-DNA interactions in greater detail, we recently developed a modified optical tweezers assay that we first applied to study T4 motor complexes containing only the TerL subunit^47^. In this study, we stalled packaging by removing ATP and measured the motor-DNA interactions in different solution conditions. In the absence of nucleotide there is almost no gripping, and the DNA slips out very rapidly (∼2,000 bp/s). In contrast, when ATP-γS is bound the motor grips DNA very persistently. Slips, where the motor temporarily releases its grip, only occur occasionally. Notably, however, when slips do occur, the transient slipping velocity is much slower, on average, than in the absence of nucleotide. Thus, nucleotide binding not only induces DNA gripping, but also regulates motor-DNA interactions during slipping. In the presence of ADP, we observed frequent transitions between gripping and slipping and highly variable slipping velocities, indicating that ADP-motor complexes have widely variable DNA interactions.

Here, we extend this measurement technique to study phage lambda motor complexes containing both the TerL and TerS subunits (the holoenzyme). The results suggest that TerS plays a previously unrecognized role in regulating motor-DNA interactions during viral DNA packaging and suggest a “sliding clamp” role for the subunit in the translocating motor. In addition to probing the role of TerS in the packaging motor, these studies provide insight into the “unit genome packaging” subtype of viruses.

## RESULTS

### Measurement Technique

To probe the dynamics of motor-DNA interactions as a function of specific bound nucleotides, we combine manipulation of single DNA molecules with optical tweezers and rapid solution exchange^47^. Phage lambda packaging is first initiated in solution with empty procapsids, recombinant lambda terminase, dsDNA, and ATP. After ∼1 minute, ATP-γS is added to produce a “stalled” procapsid-motor-DNA complex. These complexes are then manipulated with optical tweezers inside a microfluidic chamber and brought near a microcapillary tube that dispenses ATP to restart packaging (“Packaging zone”, **Figure 2**). A small stretching force (5 pN) is applied to the unpackaged DNA segment so its length versus time can be measured. Packaging is then tracked until a small fraction (∼1-6 kbp) of the 48.5 kbp lambda genome length is packaged. With this length of DNA packaged there is negligible internal force resisting DNA translocation^8, 11, 35^. To probe the motor-DNA interactions in defined conditions, the complex is then rapidly moved out of the ATP solution into a different region (the “DNA grip measurement zone”) containing either: (i) no nucleotide, (ii) poorly hydrolyzed ATP analogs (ATP*) or (iii) ADP (**Figure 2**). No further packaging is measured in the absence of ATP, so the DNA is either being gripped by terminase or slipping out of the capsid.

**Fig. 2.**
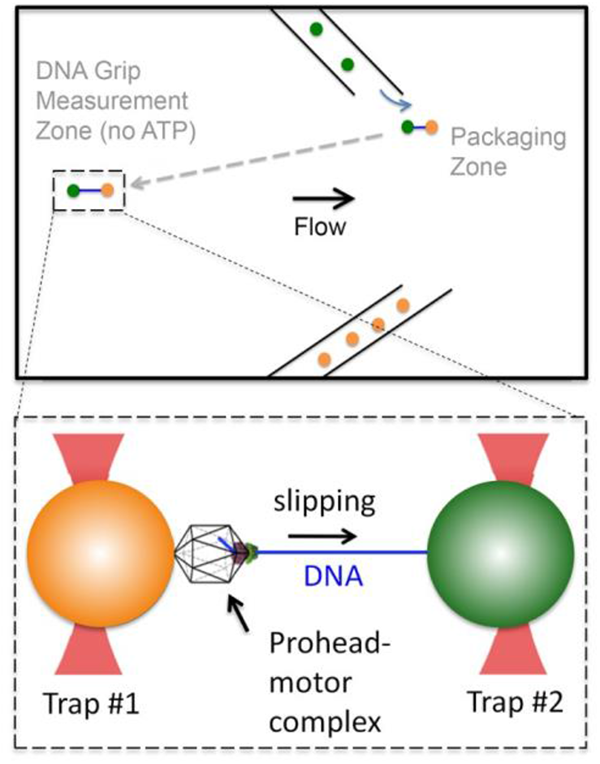
Lambda packaging complexes stalled with ATP-γS are attached by the DNA end to a microsphere (orange) and trapped with optical tweezers. This microsphere is brought near a second trapped microsphere coated with antibodies that bind the capsid (green). When exposed to ATP the motor packages the DNA. After ∼1.5-6 kbp of DNA (blue) is packaged, the complex is moved (dashed arrow) into a region containing either no nucleotide or 0.5 mM ATP-γS, AMP-PNP, or ADP. The length of the DNA outside the capsid versus time is measured.

The indicated nucleotides are added at sufficiently high concentration that they will bind rapidly to the motor subunits. Specifically, we used a 0.5 mM concentration based on the observation that 0.5 mM ATP maximizes the DNA translocation velocity. Motor grip/slip dynamics are then measured for many complexes and the ensemble of data evaluated to determine: (i) the average DNA exit velocity (including slipping and gripping events), (ii) the transient slipping velocities (the velocity of DNA movement during slipping events, excluding gripping events), (iii) the percentage time DNA is gripped by the motor, (iv) the frequency of gripping events, and (v) the duration of gripping and slipping events.

### Dependence of Motor-DNA Interactions on Nucleotides

Examples of individual measurements are shown in **Figure 3A**. Dynamic transitions between DNA gripping and slipping are observed under all conditions, but there are significant differences depending on nucleotides (**Table 1**). In the absence of nucleotide (when motor subunits are in the apo state), the motor is in a gripping conformation 44% of the time, whereas with ATP* it is gripping almost continually (99.6% of the time with AMP-PNP). Thus, ATP* binding (and presumably ATP binding) nearly always puts motor subunits in a state that allows them to persistently grip DNA. When we add ADP, in contrast, the motor grips somewhat persistently (84% of the time), but notably much less than with ATP*. The overall effect of varying nucleotide grip states on DNA retention can be inferred from measuring the average DNA exit velocities, which are 59 bp/s, 0.5-1.7 bp/s, and 11 bp/s for the apo-, ATP*- and ADP conditions, respectively (**Table 1**). These metrics indicate that interactions between motor and DNA are strongest with ATP*, weaker with ADP, and weakest with no nucleotide.

**Fig. 3.**
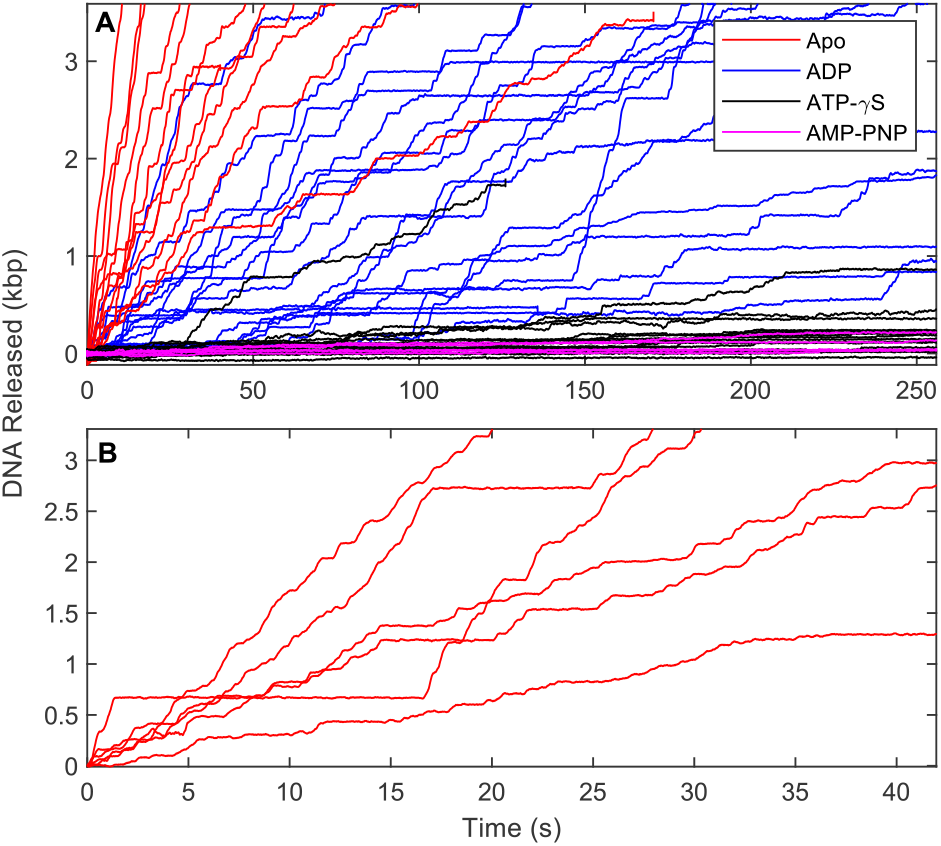
**(A)** Representative measurements of DNA length released vs. time in a solution with either no nucleotide (“apo” state, red lines), ADP (0.5 mM, blue), ATP-γS (0.5 mM, black), or AMP-PNP (0.5 mM, magenta). Y-axis indicates the length of DNA released from the nucleocapsid bound to terminase. Gripping is evidenced by horizontal portions of each curve, while slipping is evident during the sloped regions. **(B)** Zoomed plot showing examples of the pauses with no nucleotide.

**Table 1.** Average metrics characterizing the gripping/slipping dynamics, comparing lambda with T4 results measured previously^47^. Due to the low slipping velocity with the ATP analogs, we could not reliably score individual gripping events, so frequencies and durations are not reported. In the T4 study periods where slipping occurred were determined using the 2σ criterion (see methods) so the same criteria was used to analyze the lambda data presented in this table. All nucleotide conditions were 0.5 mM.

|  | % Time Gripped | Average DNA exit velocity (bp/s) | Transient slip velocity (bp/s) | Grip duration (s) | Slip duration (s) | Grip frequency (#/kbp) | Number of pull-out events |
| --- | --- | --- | --- | --- | --- | --- | --- |
| **Lambda (5 pN force)** |  |  |  |  |  |  |  |
| Apo | 44 (2.2) | 58.75 (0.4) | 120.2 (10.1) | 2.07 (0.51) | 1.95 (0.29) | 4.44 (0.37) | 53 |
| ADP | 83 (1.3) | 11 (0.08) | 76.5 (8.5) | 4.8 (1.43) | 0.86 (0.12) | 17 (1.5) | 51 |
| ATP-γS | 97 (1.2) | 1.7 (0.03) | 45.8 (3.4) | | | | 23 |
| AMP-PNP | 99.6 (0.13) | 0.47 (0.02) | 39.5 (2.3) |  |  |  | 23 |
| **T4** |  |  |  |  |  |  |  |
| Apo | 2 | 1960 (75) | 2000 (75) | 0.08 (0.02) |  |  | 25 |
| ADP | 4.6 | 844 (93) | 885 (90) | 0.28 | 1.64 |  | 35 |
| ATP-γS | 84 | 2.3 (0.54) | 33.8 (6.3) | | | | 9 |

DNA gripping by the lambda motor in the absence of nucleotide, when all motor subunits are in the apo state, was unexpected as virtually no gripping was observed in this condition with the phage T4 TerL motor^47^. We analyzed these events in detail to determine the frequency of and durations of gripping events (**Table 1**). The average duration (*i.e.*, lifetime) of a gripping event was 2.07 s while the average slip duration (*i.e.,* time required for regripping) was 1.95 s. Another notable feature is the effect of the non-gripping motor on DNA exit. The transient slipping velocity (**Figure 4** and **Table 1**) with a 5 pN force measured 120 bp/s, on average, which is ∼17-fold lower than that observed with T4 and indicates there is much stronger friction between the motor and DNA even in the apo state. There is also significant variation in the slipping velocities (standard deviation of 73.5 bp/s; see **Figure 4**), which indicates that the strength of the friction fluctuates, and the range of fluctuations is mediated by nucleotide state.

**Fig. 4.**
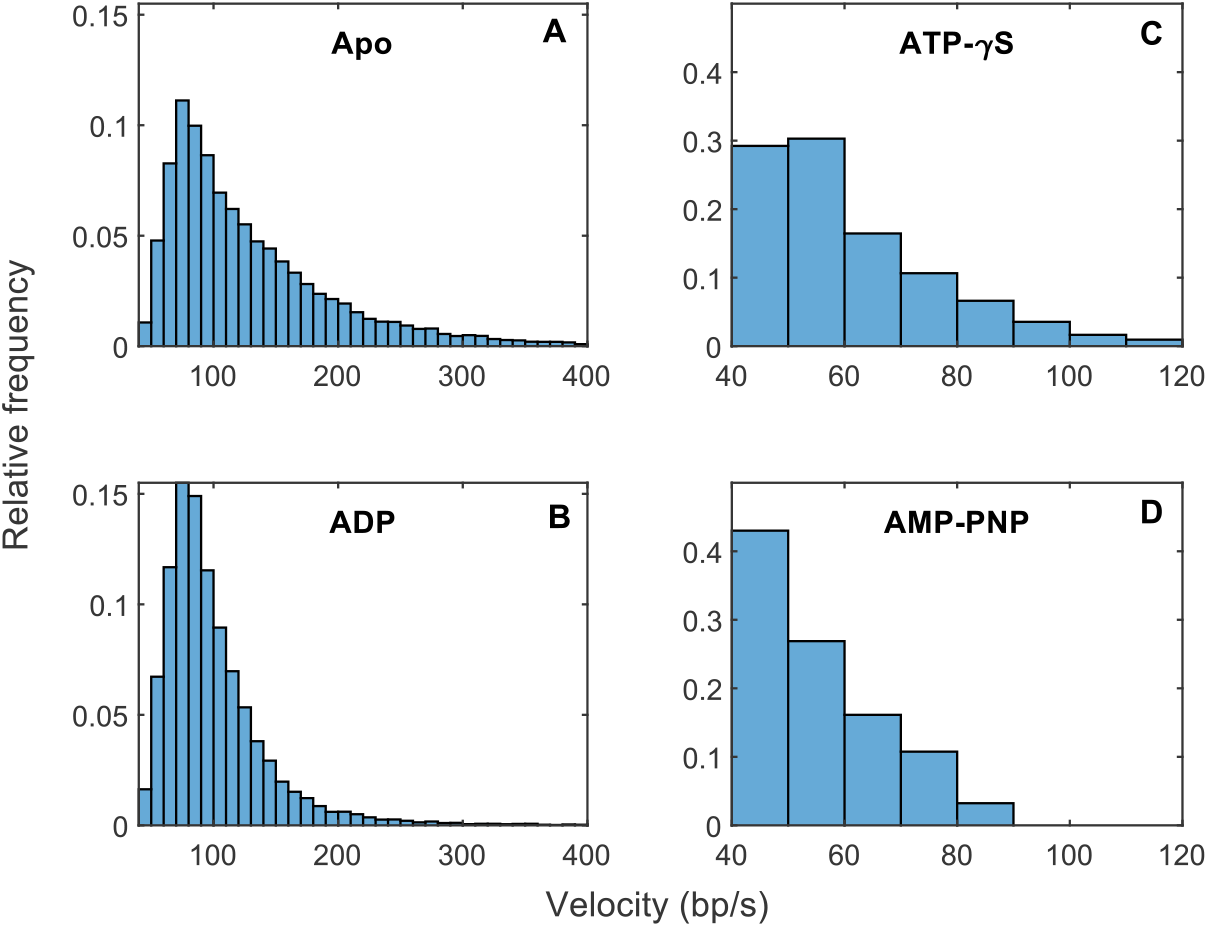
Transient slipping velocities measured in 1 s time intervals with 5 pN applied force. For these plots, and the subsequent figures, periods of slipping vs. gripping were determined using the 3σ criterion (see SI methods).

In contrast, addition of ATP* results in almost continuous gripping (**Table 1**). In our previous studies with T4 we tested only one ATP analog, ATP-γS. Here we tested both ATP-γS and AMP-PNP and although both induced highly persistent gripping, a difference was observed (Supplemental Information (SI) **Figure. S1**). With ATP-γS the DNA is gripped 97% of the time, whereas with AMP-PNP it is 99.6%. Thus, binding of AMP-PNP locks the motor in a nearly static grip. Although rare, slips are still observed with bound ATP* and it is notable that the average transient slipping velocity is ∼3-fold slower than that measured for the apo-motor (**Table 1**). Thus, binding of ATP* to the motor increases the strength of motor-DNA interactions in terms of both persistence of grip *and* friction.

Investigating the effect of bound ADP on motor-DNA interactions is of interest because, during packaging, the translocating TerL subunits have ADP transiently bound after each cycle of ATP hydrolysis. Our measurements show that with ADP bound DNA is gripped 84% of the time, ∼2x as often as in the apo condition. The frequency of these gripping events is ∼4x greater than with the apo-motor, and the average duration is ∼2.3x longer. Thus, binding of ADP also favors transition of motor subunits to a gripping state, although this state is shorter lived than with ATP*. During slipping the transient velocity with ADP (77 bp/s) is notably higher than with ATP* (40 bp/s), but again significantly lower than with no nucleotide (120 bp/s). As with ATP* this implies that motor subunits in the ADP-bound state not only grip the DNA more frequently than in the apo state, but also exert a higher frictional force on DNA when not gripping. Again, this velocity is quite variable (standard deviation of 60 bp/s, see **Figure 4**), indicating that the strength of the friction fluctuates.

### Dependence of Motor-DNA Interactions on Applied Force

The movement of DNA during slipping is overdamped motion on the timescale of the measurement (*i.e.*, there is no transient acceleration). This implies that there is a resisting friction force equal in magnitude to the force we apply with the optical tweezers (5 pN), which allows us to infer the friction force. This friction is dominated by the motor-DNA interactions since hydrodynamic drag forces on the DNA both inside and outside the capsid are negligible in comparison (calculations presented in the SI). When a given force is applied and slower slipping is observed this implies a higher friction coefficient between motor and DNA, which implies stronger motor-DNA interactions. Thus, measurements of the transient slipping velocity inform us about the nature of the friction and strength of these interactions.

As expected, we find that the transient slipping velocity increases with increasing applied force (**Figure 5A, Table S1**). It is notable that the velocity is not directly proportional to force as would be expected if the friction were dominated by hydrodynamic drag^48^. In the absence of nucleotide (apo enzyme), when the force is increased from 2 to 5 pN (a 2.5-fold increase), the velocity increases by only 1.3-fold. In contrast, increasing the force from 5 to 10 pN (a 2-fold increase) increases the velocity 1.6-fold. Similarly, in the presence of ADP, increasing the applied force from 2 to 5 pN increases velocity by only 3%, and going from 5 to 10 pN increases it by 1.3-fold (**Figure 5A**). This non-linear dependence of velocity on force is consistent with the friction being dominated by inter-molecular sliding friction^49^ between the DNA and motor, rather than hydrodynamic drag. The observed trend that there is a weaker force dependence when the friction is stronger (with ADP vs. with no nucleotide) also supports the conclusion that the friction is dominated by sliding friction^49^. The stronger friction detected with ADP implies closer contact between motor and DNA in this condition.

**Fig. 5.**
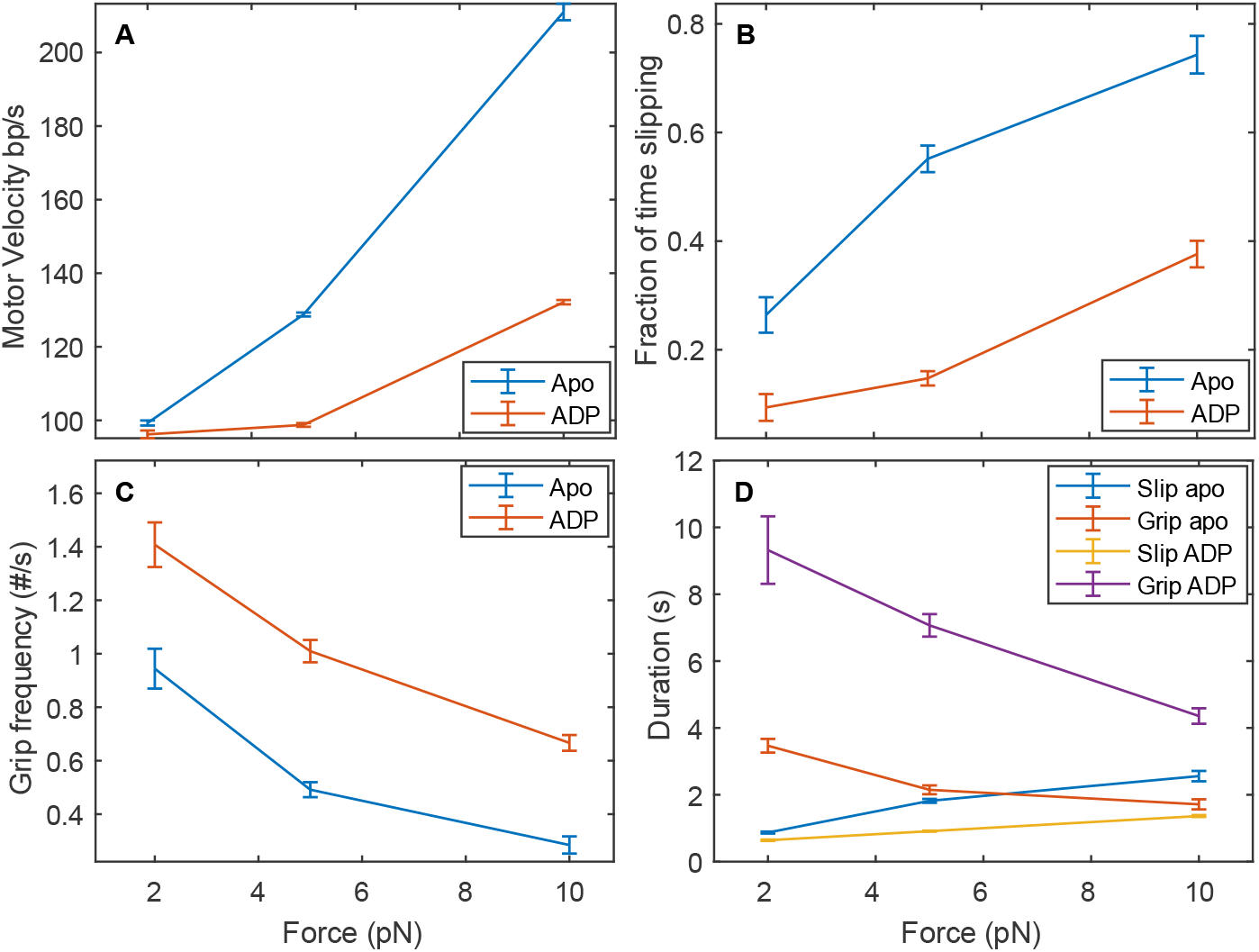
**(A)** Average transient slipping velocity vs. applied force. **(B)** Gripping frequency vs. force, calculated as the number of pauses per amount of time spent slipping. Blue points indicate measurements with no nucleotide and orange points those with 0.5 mM ADP. Error bars indicate standard errors of the means. **(C)** Mean grip and slip durations vs. force. Blue: slipping with no nucleotide, orange: gripping with no nucleotide, yellow: slipping with 0.5 mM ADP, and purple: gripping with 0.5 mM ADP. **(D)** Fraction of time spent slipping vs. force, calculated as fraction of 1 s time bins in which slipping occurs, with no nucleotide (blue) or 0.5 mM ADP (orange). Error bars indicate standard errors of the means.

In both the apo and ADP conditions, the percentage of time spent slipping increases with increasing force (**Figure 5B, Table S1**). Increasing force is expected to lead to accelerated rupture of the motor’s grip on DNA and to hinder re-gripping since it produces higher slipping velocity. Consistent with these expectations we find that both the frequency and average duration of gripping events decrease with increasing force (**Figure 5C&D, Table S1**). It is notable that in the apo condition increasing force from 2 to 10 pN has a stronger influence on gripping frequency (decreasing it 3.4-fold) than on average grip duration (decreasing it 2-fold), whereas with ADP both gripping frequency and average grip duration are reduced 2.1-fold. This is consistent with re-gripping being more strongly hindered in the apo state due to the higher transient slipping velocity. Increasing force also results in a greater increase in the average slip duration for the apo condition than with ADP (**Figure 5C**), which further supports this conclusion.

### DNA End Clamp Mechanism

Significant slipping occurs during ATP-powered packaging, and increased slipping occurs when the ATP concentration is lowered^39^. If unchecked, slipping during the early stages of packaging might allow the partially packed DNA to entirely slip out and dissociate from the capsid. In our prior studies with T4 we discovered a remarkable DNA “end clamp” mechanism that prevents this^47^. Here we show that phage lambda exhibits the same mechanism. Independent of the length of DNA initially packaged, slipping suddenly arrests when the motor approaches the end of the packaged DNA. The length of DNA left hanging out of the capsid is consistent with the packageable length of the substrate DNA (∼10 kbp) suggesting that the motor is stably bound at or near the duplex end (**Figure 6B**), with the measured variations attributable to limits of the accuracy of absolute length measurements with the optical tweezers instrument. End clamping occurs in all the solution conditions studied, showing that it is independent of the nucleotide binding states of motor subunits, and the end-clamped complex is stable for several minutes even with a 5 pN applied force.

**Fig. 6.**
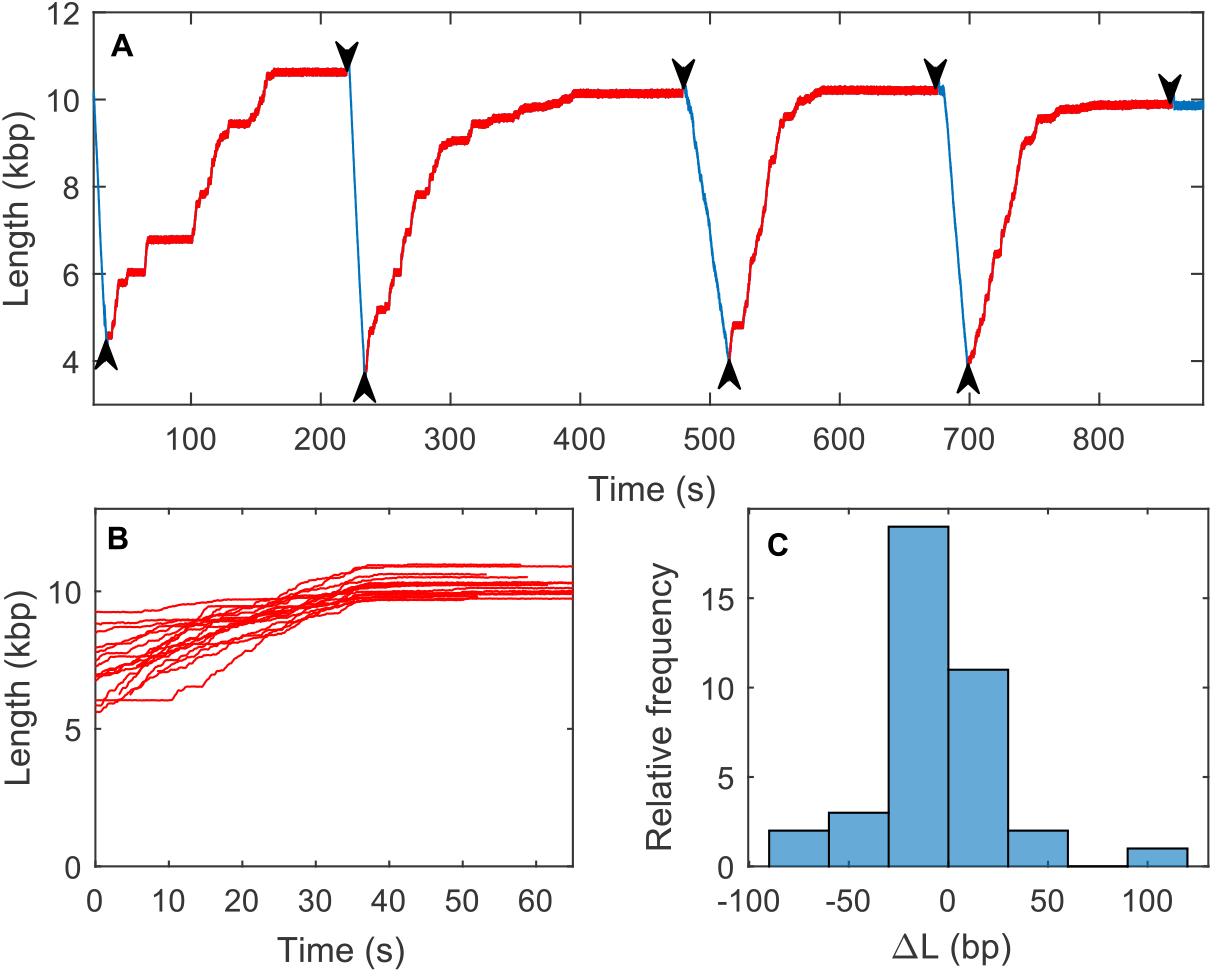
**(A)** Example of repeated slip/clamp/re-package measurements with 10 pN force. The DNA is packaged with ATP (blue) and when the solution is changed from ATP to ADP (upward arrows) the DNA slips out but is end clamped (red). Each time the complex is moved back into ATP (downward arrows) the DNA is re-packaged. **(B)** Examples of length of DNA released vs. time measured during slipping with no nucleotide and 5 pN force. Slipping usually arrests when the measured length is around 10 kb, consistent with the full length of the DNA substrate. The observed variation in the final lengths is fully attributable to limits in the accuracy of absolute DNA length measurements with the optical tweezers due variations in the sizes of microspheres used (see SI). **(C)** Histogram of differences in the DNA length measured in repeated end clamp events with single complexes.

From this end-clamped state, we find that packaging can be restarted by performing exchange back into a solution containing ATP. Packaging, slipping, end clamping, and re-packaging can be repeated multiple times and no differences in the packaging or gripping/slipping dynamics are observed (**Figure 6A**). Measurements of the length differences following repeated clamping events with single complexes also allows a more precise determination of the repeatability of the position where the DNA is held. The standard deviation of the repeated clamp position was only 32 bp (**Figure 6C**), which is consistent with resolution of the measurement (when moving to different positions in the flow chamber), and therefore implies that the DNA is being clamped exactly at, or very near to its end.

## DISCUSSION

The phage lambda motor has broad similarity to all characterized viral genome packaging motors, including T4, but also important differences which may be related to differences in their function^2, 4^. Specifically, phage T4 represents the headful packaging class of viruses while lambda represents a distinct major class that instead packages a unit-length genome (**Figure 1**). This has implications for how the translocating motor recognizes the genome end and cuts the duplex to terminate packaging. Moreover, while all terminase enzymes employ a TerS subunit that is essential for initiation of packaging *in vivo*, whether this subunit plays any role during DNA translocation has remained unclear. I*n vitro* T4 packaging studies have employed motor complexes assembled without TerS, which importantly showed that TerL is sufficient to translocate DNA but could not investigate potential roles of TerS to be probed^28^. In the present work, lambda provides a system where the terminase can be assembled *in vitro* into a functional packaging motor containing both subunits in the hetero-oligomeric complex (TerL•TerS) ^23–25^.

Our present studies, when compared to our findings with T4, reveal universal features of the terminase motors, provides fundamental information on a second major class of packaging motors, and suggests a role for the conserved TerS subunit during DNA packaging. In both motors, we find that the persistence of gripping is strongest with ATP* bound, weaker with ADP bound, and weakest with no nucleotide (apo terminase). These findings provide support for a conserved universal mechanism for terminase motor function in which ATP binding induces motor subunits to undergo a conformational change that causes them to grip DNA tightly^33, 41, 44^. Hydrolysis of the bound nucleotide and release of P_i_ affords a bound ADP state with weakened grip, and subsequent ADP release (apo conformation) releases the grip. However, our measurements show that there is additional regulation of motor-DNA interactions beyond just DNA grip/release. Namely, the transient slipping velocity when the DNA is not gripped is strongly affected by nucleotide binding. It is slowest with ATP* bound, faster with ADP bound, and fastest with no nucleotide. This implies that there is sliding friction between the motor and DNA that is modulated by nucleotide binding to non-gripping terminase subunits. This friction, although it may impose additional load on the motor if it acts during translocation, also conveys a benefit by limiting exit of the DNA from the capsid when the motor loses grip.

The qualitative similarity in these characteristics, *i.e.,* nucleotide dependent DNA gripping and friction, for these two classes of terminase motors suggests they are conserved universal features. On a *quantitative* level, however, we observe striking differences between the lambda motor assembled from both requisite TerL and TerS subunits and the T4 motor containing only TerL. The measured motor-DNA interactions are, in every condition, much stronger for the lambda holoenzyme (**Table 1**). With ATP-γS gripping is persistent in both cases, but the percent time slipping is 5-fold lower for lambda. With lambda we also tested AMP-PMP and found it to induce almost 100% persistent gripping and ∼4x lower average DNA exit velocity than ATP-γS (**Table 1, Figure S1**). This difference is likely attributable to faster dissociation and/or hydrolysis/product release of ATP-γS, which indicates that AMP-PNP is a better analog for freezing terminase in an ATP bound-like conformation.

In further comparison to T4, the percent time gripped with ADP is 18-fold higher and the average duration of gripping events 17-fold longer for lambda. This results in a remarkable 77-fold lower average DNA exit velocity. But perhaps the most striking difference, not observed with T4, is that frequent gripping and highly slowed slipping is observed with lambda even without nucleotides, resulting in a 33-fold lower average DNA exit velocity. This indicates that the lambda motor, even in the apo state, frequently fluctuates into a conformation that enables it to grip DNA.

These large differences between lambda and T4 are likely attributable to the presence of TerS in the lambda motor. The stronger motor-DNA interactions could be due to TerS directly interacting with the DNA, as it does during initiation of packaging^2^, or to an influence of TerS on the conformation of TerL, since TerS is directly bound to TerL in the holoenzyme. Within this context, structural studies on lambda terminase holoenzyme reveal that TerS forms a decameric ring of sufficient size to encircle DNA^25^. The five TerL subunits extend from the ring in a starfish-like geometry, but likely clamp down in the assembled motor to simultaneously interact with the DNA and bind to the portal ring of the procapsid. Based on these observations, we suggest that the TerL ring serves as the *bona fide* motor while the decameric TerS ring serves as a sliding clamp, analogous to those involved in DNA replication that ensures high processivity^50, 51^, such that an entire genome is packaged in a single DNA binding event. This is consistent with our finding that during active packaging with saturating ATP the lambda motor exhibited ∼5-fold fewer detected slips per length of DNA translocated than the T4 motor^34, 35^.

Our finding that the lambda motor frequently grips DNA even in the absence of nucleotide also suggests an explanation for a previous observation that long pauses that significantly affect the packaging kinetics frequently occur during ATP-fueled DNA translocation^39^. Notably, these pauses were found to have an average duration of 2.0 s, which is almost the same as the average duration of the gripping events we measure here with no nucleotide (2.1 s). This suggests that the pauses during packaging are due to stochastic fluctuation of motor subunits in the apo state into a DNA gripping conformation. The long duration of these pauses indicates this is an off-pathway transition with respect to the main ATP hydrolysis cycle, since during normal translocation with saturating ATP each hydrolysis-DNA translocation event occurs in less than 0.01 seconds^41^. This capability of the motor to pause and then resume packaging may convey a benefit during packaging by helping to arrest slipping whenever the motor temporarily loses grip. Lambda phage DNA is subject to recombination events that can produce DNA-DNA junctions that could impede translocation^52, 53^. The ability of the motor to pause and idle without hydrolyzing ATP when it encounters a DNA junction and then later resume packaging may be an important feature.

The kinetic model we previously proposed for the DNA grip/slip dynamics observed with T4^47^ can be modified to account for the differences we observe with lambda. This would require adding a pathway that allows fluctuation of motor subunits in the apo state into a conformation that can either grip DNA or exert high friction. The “minimum friction” state proposed to explain the much higher T4 slipping velocity in the apo and ADP bound states would need to be eliminated. It should be noted, however, that this model was based on a common previous assumption that only one motor subunit grips DNA at a time^21, 33, 36, 37, 39^. Recent structural studies of the phage phi29 motor suggest that this might not be the case. Specifically, it was found that the phage phi29 motor can adopt either a planar ring or a helical split-ring conformation^54^, and suggested a model in which multiple ATP-bound subunits can grip simultaneously^55^. Although the phi29 motor is not a terminase and lacks a TerS subunit, its ATPase has significant homology with TerL. If a similar mechanism applies to the lambda and T4 terminase motors, then previously proposed models would need to be revised. In particular, it was proposed that transient ATP* or ADP dissociation from one subunit causes onset of slipping^47^. An alternative model compatible with multi-subunit gripping is that slipping is initiated by force-induced rupture of grip while one or more subunits are in a grip-competent conformation. Within this model the slow slipping observed with ATP* bound is attributable to subunits in this conformation exerting high friction, and regripping attributable to stochastic recapture of the DNA while subunits remain in this conformation.

Another general observation is that both lambda and T4 both exhibit a DNA end clamp, preventing DNA uncoupling from the motor-capsid even in the absence of nucleotides, suggesting that this is a conserved property of terminase motors. This mechanism increases the efficiency of packaging since it ensures initiation of packaging is essentially irreversible once started. Without such a mechanism, duplex release from the capsid could occur if the motor slipped when only a small length of DNA was packaged, and the entire complex would then need to be reassembled. This mechanism is especially advantageous for lambda, since assembly on the initiation site by the holoenzyme necessitates a cleavage event, meaning loss of the duplex would be irrevocable. Moreover, a dissociated duplex would render the exposed genome end to host cell exonucleases which would degrade unprotected DNA^56^.

The molecular mechanism by which the end clamp works remains unclear, but based on the principle of microscopic reversibility, we suggest that the end-clamped complex in phage lambda most likely recapitulates the initiation complex assembled at the *cos* site on the DNA when terminase first brings the genome end to the portal nanochannel in the procapsid shell^2, 4^. Importantly, our finding that packaging, slipping, end clamping, and re-packaging can be repeated multiple times and no differences in the packaging or gripping/slipping dynamics are observed suggests that the transition from the initiation complex to the translocating motor complex is a fully reversible transition. In future studies our measurement technique could be further extended to probe the stability of the end-clamped state (likely equivalent to the packaging initiation state) as a function of time, applied force, solution conditions, and *cos* sequence variation.

## CONCLUSIONS

In summary, we show that the phage lambda motor, qualitatively like that of phage T4, exhibits nucleotide-dependent DNA gripping and friction between motor and DNA. ATP analog binding induces highly persistent DNA gripping and high friction between motor and DNA that limits slipping, whereas ADP induces intermittent gripping and weaker friction. Thus, our findings provide evidence for several conserved features of terminase viral packaging motors. A striking difference, however, is that the lambda motor exhibits dramatically stronger DNA interactions than the T4 TerL motor, both in terms of persistence of gripping and friction. Moreover, the lambda motor frequently grips DNA even in the absence of nucleotide. These differences are likely due to the inclusion of the small terminase subunit (TerS) in the lambda motor complexes. Our findings suggest that TerS plays a previously unrecognized and important role to increase the motor processivity by acting as a “sliding clamp”. We also show that both lambda and T4 also have a nucleotide-independent DNA end-clamping mechanism that increases the efficiency of packaging. In the unit genome packaging class of viruses such as phage lambda the end clamp state is likely equivalent to the packaging initiation complex, such that our method provides an avenue to study factors affecting the stability of the initiation complex in individual packaging-competent motors.

## MATERIALS AND METHODS

Optical tweezer single-molecule experiments were performed at varying applied forces in a buffer solution containing 25 mM Tris-HCl pH 7.5, 5 mM MgCl2, and varying nucleotides at 0.5 mM, as described in the text. Complexes were assembled using purified procapsids, purified terminase proteins, cos-containing DNA, and integration host factor (IHF) from *E. coli,* and then prestalled. Slipping and gripping events were detected based on velocity threshold analyses. More details on the methods of protein purification, substrate and complex production, optical tweezers measurements, and data analysis are provided in the SI.

## ACKNOWLEDGEMENTS

This work was supported by NIH grants R01GM118817 (DES) and R01GM127365 (CEC) and T32 GM008326 (to BR through the Molecular Biophysics Training Program at the University of California, San Diego).

## REFERENCES

1 SR Casjens, The DNA-packaging nanomotor of tailed bacteriophages. Nat. Rev. Microbiol. 9, 647–657 (2011).

2 VB Rao, M Feiss, Mechanisms of DNA packaging by large double-stranded DNA viruses. Annu. review Virology 2, 351–378 (2015).

3 C Hetherington, J Moffitt, P Jardine, C Bustamante, Viral DNA packaging motors in Comprehensive Biophysics. (Elsevier Inc.), pp. 420–446 (2012).

4 CE Catalano, MC Morais, Viral genome packaging machines: Structure and enzymology. The Enzym. 50, 369–413 (2021).

5 B Roizman, Herpes simplex viruses. Fields Virology pp. 2501–2602 (2007).

6 RL Calendar, ST Abedon, The bacteriophages. (Oxford University Press on Demand) Vol. 2, (2006).

7 SC Riemer, VA Bloomfield, Packaging of DNA in bacteriophage heads: some considerations on energetics. Biopolym. Orig. Res. on Biomol. 17, 785–794 (1978).

8 DE Smith, et al., The bacteriophage φ29 portal motor can package DNA against a large internal force. Nature 413, 748–752 (2001).

9 J Kindt, S Tzlil, A Ben-Shaul, WM Gelbart, DNA packaging and ejection forces in bacteriophage. Proc. Natl. Acad. Sci. 98, 13671–13674 (2001).

10. ^10^ DE Smith, Single-molecule studies of viral DNA packaging. Curr. opinion Virology 1, 134–141 (2011).

11. ^11^ A Evilevitch, L Lavelle, CM Knobler, E Raspaud, WM Gelbart, Osmotic pressure inhibition of DNA ejection from phage. Proc. Natl. Acad. Sci. 100, 9292–9295 (2003).

12. ^12^ PK Purohit, J Kondev, R Phillips, Mechanics of DNA packaging in viruses. Proc. Natl. Acad. Sci. 100, 3173–3178 (2003).

13. ^13^ AS Petrov, SC Harvey, Packaging double-helical DNA into viral capsids: structures, forces, and energetics. Biophys. Journal 95, 497–502 (2008).

14. ^14^ AY Lyubimov, M Strycharska, JM Berger, The nuts and bolts of ring-translocase structure and mechanism. Curr. opinion structural biology 21, 240–248 (2011).

15. ^15^ JF Allemand, B Maier, DE Smith, Molecular motors for DNA translocation in prokaryotes. Curr. opinion biotechnology 23, 503–509 (2012).

16. ^16^ S Liu, G Chistol, C Bustamante, Mechanical operation and intersubunit coordination of ring-shaped molecular motors: insights from single-molecule studies. Biophys. Journal 106, 1844–1858 (2014).

17. ^17^ JA Hayes, BA Kelch, DNA packaging: The translocation motor in Encyclopedia of Virology (Fourth Edition), eds. DH Bamford, M Zuckerman. (Academic Press, Oxford), Fourth edition, pp. 148–159 (2021).

18. ^18^ RK Lokareddy, CFD Hou, F Li, R Yang, G Cingolani, Viral small terminase: A divergent structural framework for a conserved biological function. Viruses 14, 2215 (2022).

19. ^19^ AS Al-Zahrani, et al., The small terminase, gp16, of bacteriophage T4 is a regulator of the DNA packaging motor. J. biological chemistry 284, 24490–24500 (2009).

20. ^20^ BJ Hilbert, et al., Structure and mechanism of the ATPase that powers viral genome packaging. 2Proc. Natl. Acad. Sci. 112, E3792–E3799 (2015).

21. ^21^ S Sun, et al., The structure of the phage T4 DNA packaging motor suggests a mechanism dependent on electrostatic forces. Cell 135, 1251–1262 (2008).

22. ^22^ H Mao, et al., Structural and molecular basis for coordination in a viral DNA packaging motor. Cell reports 14, 2017–2029 (2016).

23. ^23^ NK Maluf, Q Yang, CE Catalano, Self-association properties of the bacteriophage λ terminase holoenzyme: implications for the DNA packaging motor. J. Molecular Biology 347, 523–542 (2005).

24. ^24^ NK Maluf, H Gaussier, E Bogner, M Feiss, CE Catalano, Assembly of bacteriophage lambda terminase into a viral DNA maturation and packaging machine. Biochemistry 45, 15259–15268 (2006)

25. ^25^ NS Prokhorov, et al., Biophysical and structural characterization of a viral genome packaging motor. bioRxiv pp. 2022–09 (2022).

26. ^26^ TC Yang, D Ortiz, GC Lander, CE Catalano, et al, ., Thermodynamic interrogation of the assembly of a viral genome packaging motor complex. Biophys. Journal 109, 1663–1675 (2015).

27 A Roy, A Bhardwaj, P Datta, GC Lander, G Cingolani, Small terminase couples viral DNA binding to genome-packaging ATPase activity. Structure 20, 1403–1413 (2012).

28 S Grimes, PJ Jardine, D Anderson, Bacteriophage φ29 DNA packaging. Adv Virus Res 58, 255–294 (2002).

29 CE Catalano, M Feiss, CE Catalano, Bacteriophage lambda terminase and the mechanism of viral DNA packaging. Viral Genome Packag. Mach. Genet. Struct. Mech. pp. 5–39 (2005).

30 KR Kondabagil, Z Zhang, VB Rao, The DNA translocating ATPase of bacteriophage T4 packaging motor. J. Molecular Biology 363, 786–799 (2006).

31 OW Bayfield, et al., Cryo-em structure and in vitro DNA packaging of a thermophilic virus with supersized t=7 capsids. Proc. Natl. Acad. Sci. 116, 3556–3561 (2019).

32 HK Fung, et al., Structural basis of DNA packaging by a ring-type ATPase from an archetypal viral system. Nucleic acids research 50, 8719–8732 (2022).

33 YR Chemla, et al., Mechanism of force generation of a viral DNA packaging motor. Cell 122, 683–692 (2005).

34 DN Fuller, DM Raymer, VI Kottadiel, VB Rao, DE Smith, Single phage T4 DNA packaging motors exhibit large force generation, high velocity, and dynamic variability. Proc. Natl. Acad. Sci. 104, 16868–16873 (2007).

35 DN Fuller, et al., Measurements of single DNA molecule packaging dynamics in bacteriophage λ reveal high forces, high motor processivity, and capsid transformations. J. Molecular Biology 373, 1113–1122 (2007).

36 JR Moffitt, et al., Intersubunit coordination in a homomeric ring ATPase. Nature 457, 446–450 (2009).

37 G Chistol, et al., High degree of coordination and division of labor among subunits in a homomeric ring ATPase. Cell 151, 1017–1028 (2012).

38 S Liu, et al., A viral packaging motor varies its DNA rotation and step size to preserve subunit coordination as the capsid fills. Cell 157, 702–713 (2014).

39 JM Tsay, J Sippy, M Feiss, DE Smith, The q motif of a viral packaging motor governs its force generation and communicates ATP recognition to DNA interaction. Proc. Natl. Acad. Sci. 106, 14355–14360 (2009).

40 JM Tsay, et al., Mutations altering a structurally conserved loop-helix-loop region of a viral packaging motor change DNA translocation velocity and processivity. J. Biol. Chem. 285, 24282–24289 (2010).

41 D Ortiz, et al., Walker-a motif acts to coordinate ATP hydrolysis with motor output in viral DNA packaging. J. Molecular Biology 428, 2709–2729 (2016).

42 D Ortiz, et al., Functional dissection of a viral DNA packaging machine’s Walker B motif. J. Molecular Biology 431, 4455–4474 (2019).

43 D Ortiz, et al., Evidence that a catalytic glutamate and an ‘arginine toggle’ act in concert to mediate ATP hydrolysis and mechanochemical coupling in a viral DNA packaging motor. Nucleic acids research 47, 1404–1415 (2019).

44 VI Kottadiel, VB Rao, YR Chemla, The dynamic pause-unpackaging state, an off-translocation recovery state of a DNA packaging motor from bacteriophage T4. Proc. Natl. Acad. Sci. 109, 20000–20005 (2012).

45 N Keller, DJ delToro, DE Smith, Single-molecule measurements of motor-driven viral DNA packaging in bacteriophages phi29, lambda, and T4 with optical tweezers. Mol. Mot. Methods Protoc. 1805, 393–422 (2018).

46 Q Yang, NK Maluf, CE Catalano, Packaging of a unit-length viral genome: the role of nucleotides and the gpd decoration protein in stable nucleocapsid assembly in bacteriophage λ. J. Molecular Biology 383, 1037–1048 (2008).

47 M Ordyan, I Alam, M Mahalingam, VB Rao, DE Smith, Nucleotide-dependent DNA gripping and an end-clamp mechanism regulate the bacteriophage T4 viral packaging motor. Nat. communications 9, 5434 (2018).

48 IS Gabashvili, AY Grosberg, Dynamics of double stranded DNA reptation from bacteriophage. J. Biomol. Struct. Dyn. 9, 911–920 (1992).

49 A Ward, et al., Solid friction between soft filaments. Nat. materials 14, 583–588 (2015).

50 GL Moldovan, B Pfander, S Jentsch, Pcna, the maestro of the replication fork. Cell 129, 665–679 (2007).

51 C Gaubitz, et al., Cryo-em structures reveal high-resolution mechanism of a DNA polymerase sliding clamp loader. Elife 11, e74175 (2022).

52 CR Hillyar, Genetic recombination in bacteriophage lambda. Biosci. Horizons: The Int. J. Stud. Res. 5 (2012).

53 FW Stahl, Recombination in phage λ: one geneticist’s historical perspective. Gene 223, 95–102 (1998).

54 M Woodson, et al., A viral genome packaging motor transitions between cyclic and helical symmetry to translocate dsDNA. Sci. Adv. 7, eabc1955 (2021).

55 J Pajak, et al., Atomistic basis of force generation, translocation, and coordination in a viral genome packaging motor. Nucleic acids research 49, 6474–6488 (2021).

56 N Sternberg, J Coulby, Recognition and cleavage of the bacteriophage p1 packaging site (pac): I. differential processing of the cleaved ends in vivo. J. Molecular Biology 194, 453–468 (1987).

