## Supplemental Information for "Regulation of phage lambda packaging motor-DNA interactions: Nucleotide independent and dependent gripping and friction"

### Supplementary Information for Regulation of Motor-DNA Interactions in the Hetero-oligomeric Phage Lambda Terminase Holoenzyme

**Authors:** Brandon Rawson, Mariam Ordyan, Qin Yang, Jean Sippy, Michael Feiss, Carlos E. Catalano, Douglas E. Smith

**Corresponding Authors:** Douglas E. Smith, University of California San Diego, Department of Physics, 9500 Gilman Drive, La Jolla CA 92093, (858) 736-5340,; Carlos E. Catalano, University of Colorado Anchutz Medical Campus, Department of Pharmaceutical Sciences, 4200 E. Montlake Blvd, Aurora, CO, 80045, (303) 724-0011,

#### Further Details of the Methods

##### Phage lambda procapsids

Lambda procapsids were obtained from *E. coli* lambda lysogen NS428 as described previously[1]. The strain contains a mutated gene preventing packaging, resulting in an accumulation of empty procapsids. The lysogen was grown to OD600 of 0.3 at 32 °C before being induced by moving to 45 °C for 15 minutes and then 38 °C for 60 minutes. After induction the culture is pelleted and resuspended in 0.5 ml TMS (50 mM Tris, pH 7.8, 10 mM $MgCl_2$, 100 mM NaCl) containing 40 units RNase-free DNase (Roche). The lysate was clarified by centrifuging at 10,000 rpm for 10 minutes, followed by 13,000 g for 3 minutes. The supernatant was then loaded on top of a linear 10-30% sucrose gradient in 0.5x TMS buffer (25 mM Tris, pH 7.8, 5 mM $MgCl_2$, 50 mM NaCl) and centrifuged at 40,000 rpm for 64 minutes. The band that contains the procapsids was collected and detected by observing the scattering of a light beam off the procapsids. They were then diluted in TMS buffer and pelleted at 40,000 rpm for 102 minutes. The pellets were re-suspended in a total of 200 µl TMS buffer by standing at 4 °C overnight. They were then re-banded on a second set of sucrose gradients, pelleted, and resuspended in a final volume of 100 µl and stored at 4 °C.

##### Terminase proteins

Terminase proteins were obtained from *E. coli* extracts, based on long-standing methods[2]. In brief, the *E. coli* lysogen OR1265 was grown at 32 °C for 8 hours in a 5 mL culture of TB broth and 50 µg/ml Amp before being transferred to a 50 mL TB-Amp broth to grow overnight. The overnight culture is then used in its entirety to inoculate a 1 L TB-Amp and grown to an OD600 of 0.6 before being induced by adding 1L of 62 °C TB broth. The induced cells are then grown at 45°C for 15 minutes, followed by 45 minutes at 42°C. The cells are then pelleted by centrifuging at 5000 g for 20 min before being stored at -80°C. The pellet is then resuspended in 50 mL of low imidazole His Buffer (20 mM Tris pH 8.6, 500 mM NaCl, 10% glycerol, 7 mM β-Me, 20 mM imidazole), 500 µl GE's PI, and lysozyme at 0.4 mg/ml and kept on ice for 30 minutes. The cells are then sonicated using 10 sets of 10 seconds at a 90% duty cycle before finally centrifuged at 13,000 g for 30 min. The supernatant containing the terminase proteins is then removed and stored at either -20°C for short term or -80°C for long term storage.

##### Sample preparation

Lambda procapsids and terminase proteins were prepared as described above. A 13,881 bp plasmid (pJM1) containing the λ *cos* site, which is necessary to initiate packaging, was prepared as follows. The plasmid was cut with restriction enzyme NcoI and used as a template for PCR with labeled primers: Digoxygenin-5'-TCGATAATCGTGAAGAGTCGGCGAGCCTGGTTAG-3' (forward) and Biotin-5'-TACGTCGAAGTGACCAACTAGGCGGAATCGGTAG-3' (reverse) (IDT, Inc.). DNA was generated using the LA

Taq PCR kit (Takara Bio, Inc), resulting in a 13,681 bp biotin-labeled dsDNA for use in packaging, with a ~10.1 kbp section that can be packaged after *cos* cleavage.

**Single molecule assay**

The optical tweezer instrument was calibrated by an established method using DNA molecules as metrology standards[3]. In brief, the calibration parameters are the trap compliance $\gamma$, force scale factor $\alpha$, length scale factor $\beta$, and displacement offset factor $V_0$. $\alpha$ is determined by finding the measured voltage of the DNA overstretch plateau, which occurs at a well-documented force[4]. $\beta$ is determined by measuring the trap separation voltage for two different DNA lengths and using the worm-like chain (WLC) model which tells us the DNA contour length is equal to its extension at 33.4 pN in our solution conditions. $V_0$ and $\gamma$ are then both determined by doing a dual parameter fit of the WLC model for both DNA lengths for the pre-overstretch segment.

The packaging measurements were done using a slightly modified version of established protocols in which procapsid-motor-DNA complex is first assembled in a bulk reaction and later restarted. Lambda procapsids, terminase, and pJM1 DNA are mixed in concentrations of 17 nM, 250 nM, and 11 ng, respectively, in a solution consisting of 25 mM Tris-HCl pH 7.5 and 5 mM $MgCl_2$. The mixture is incubated at room temperature for 5 minutes before adding ATP to 0.5 mM. After a 45 second incubation ATP-$\gamma$S is added to 0.5 mM to stall the reaction.

The resulting procapsid-motor-DNA complex, with the biotin labeled end of the DNA hanging out, is then mixed with 2.1 µm diameter streptavidin coated microspheres (Spherotech) to bind the labeled DNA end. This mixture is incubated on a rotating incubator at room temperature for 20 minutes. Antisera against lambda phage, containing antibodies that bind the lambda procapsid, are mixed with 2.3 µm diameter protein-G microspheres (Spherotech) at a ratio of 2.5 µl antibody to 5 µl beads (5% w/v) and incubated for 30 minutes. The main flow cell buffer, where DNA grip/slip measurements were conducted, consisted of 25 mM Tris-HCl pH 7.5, 5 mM $MgCl_2$, and 0.05 g $L^{-1}$ BSA. A separate packaging region of the chamber contained the same buffer with the addition of 0.5-1 mM ATP dispensed from a side channel to fuel packaging.

**Slipping/gripping measurements and data analysis**

The dual optical trap system uses a laser split into orthogonally polarized beams, one fixed and the other movable by a piezo-actuated mirror[3]. The deflections of the fixed beam are measured at 1 kHz by imaging the back focal plane of the objective onto a position-sensing detector, whose signal is digitized and then scaled using the calibration parameters mentioned above to determine the force. The DNA length is similarly calculated using the separation and compliance of the traps, force on the trapped bead, and force-extension relationship of DNA from the worm-like chain model[5,6]. The slipping measurements are recorded in the same way as packaging measurements using a feedback control system which varies the separation between the two traps to keep the applied force constant.

Due to inherent measurement/instrument noise we first conducted control experiments in which DNA molecules alone were tethered between two microspheres to determine the uncertainty in velocity measurements for a stationary system. For a 1 s detection window we measure and average velocity close to zero (0.004 bp/s), but with a standard deviation ($\sigma$) of 14.5 bp/s that characterizes the uncertainty due to Brownian fluctuations and instrument noise. For all the results shown in the figures, except for Table 1, the threshold velocity to identify a period of gripping was set to be $3\sigma$=43.5 bp/s. For the results presented in Table 1 the threshold was set to $2\sigma$ for direct comparison with the previous T4 studies which used this threshold[7]. For the rest of our analyses this threshold was set to $3\sigma$ because analyses comparing the total length slipped with the sum of the lengths of the individual slips showed that this provides a more accurate method for discerning periods of gripping from periods of slipping.

**Table S1.** Average metrics characterizing the lambda gripping/slipping dynamics with varying applied forces. Periods where slipping occurred were determined using the 3σ velocity threshold (see methods). Frequencies and durations were not calculated for individual grip/slip events with the ATP analogs because they could not be reliably detected due to the low transient slipping velocity in this condition. They were not calculated for the 20 pN measurements because the high slipping velocity introduces significant bias in the detection of shorter gripping events relative to the lower force measurements. Each of the listed nucleotides were added at a concentration of 0.5 mM.

| | Transient Slip Velocity (bp/s) | Average Exit Velocity (bp/s) | % Time Gripped | Grip Frequency (#/kbp) | Grip Duration (s) | Slip Duration (s) |
|---|---|---|---|---|---|---|
| **2 pN Force** | | | | | | |
| Apo | 99.3 (0.69) | 20 (0.23) | 73 (3.2) | 12.4 (1.9) | 3.47 (0.2) | 0.87 (0.03) |
| ADP | 96.2 (1.05) | 6.2 (0.11) | 91 (3.2) | 20.2 (5.1) | 9.3 (1) | 0.64 (0.02) |
| **5 pN Force** | | | | | | |
| Apo | 128.8 (0.53) | 58.8 (0.4) | 45 (2.5) | 4.23 (0.35) | 2.14 (0.13) | 1.82 (0.06) |
| ADP | 98.8 (0.51) | 11 (0.08) | 85 (2.5) | 12.3 (1.03) | 7.07 (0.34) | 0.91 (0.02) |
| ATP-γS | 61.3 (0.57) | 1.8 (0.03) | 98 (0.9) | | | |
| AMP-PNP | 55.4 (1.15) | 0.48 (0.02) | 99.7 (0.11) | | | |
| **10 pN Force** | | | | | | |
| Apo | 211 (2.2) | 118 (1.6) | 26 (3.5) | 1.74 (0.3) | 1.71 (0.15) | 2.6 (0.15) |
| ADP | 132 (0.59) | 30.7 (0.21) | 62 (3.4) | 5.71 (0.34) | 4.4 (0.23) | 1.36 (0.03) |
| **20 pN Force** | | | | | | |
| Apo | 391 (7.5) | 149 (3.5) | 26 (4.8) | | | |
| ADP | 235 (2.8) | 49 (0.59) | 57 (4.8) | | | |
| ATP-γS | 72.9 (0.41) | 8.04 (0.05) | 87 (2.4) | | | |

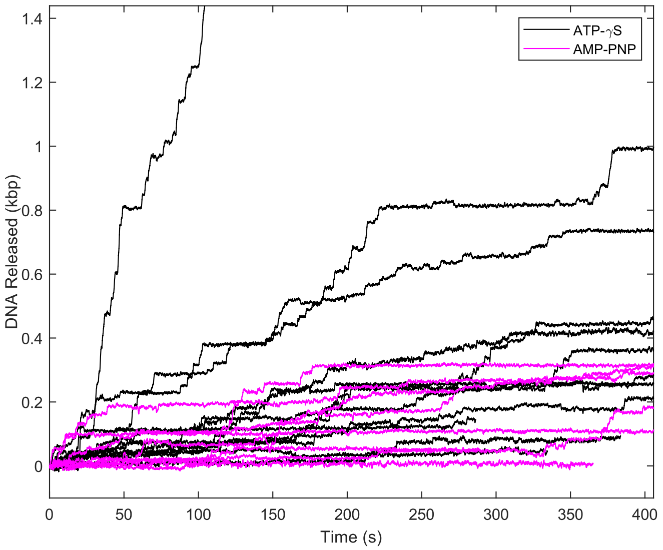


**Fig. S1.** Examples of gripping/slipping dynamics measured with ATP-γS (black lines) vs. AMP-PNP (magenta lines) with 5 pN applied force.

**Hydrodynamic Drag Calculations**

As mentioned in the text, the hydrodynamic drag forces acting on the sections of the DNA outside the capsid, inside the capsid, and threading through the portal-motor channel are all estimated to be negligible compared with the 5 pN friction force resisting DNA exit, which implies that sliding friction between the DNA and motor is the main source of friction. The finding that transient DNA slipping velocity is significantly slowed when nucleotides bind to the motor further confirms this. The hydrodynamic drag on the section of DNA outside the capsid is estimated as follows[8]. Since it is stretched to form a nearly straight line the drag is approximately that acting on a long, slender cylinder (of diameter d=2 nm and length $L_{out} = 2\ \mu m$ (at most) $F_{out} = v\ \xi_{out}$ where $v$ is the average slipping velocity (128 bp/s) and the drag coefficient is given by $\xi_{\text{out}} = \frac{2\pi L_{out}\eta}{\ln\left(\frac{L_{out}}{d}\right)}$ where $\eta$ is the solvent viscosity. These values yield a maximum drag force of only 0.11 fN.

The hydrodynamic drag on the section of DNA inside the capsid can be estimated using the Gabashvili-Grosberg theory[9]: $F_{int} = v\ \xi_{int}$, where the drag coefficient is given by $\xi_{\text{int}} = \frac{2\pi L_{int}\eta}{\ln\left(\frac{d_s - r}{r}\right)}$

where $L_{int}$ is the length of DNA inside the capsid (6 kbp at the largest), $d_s$ the average inter-axial spacing between the packed DNA segments (less than 3.2 nm for the lengths of DNA we have packaged[10]), and $r$=1 nm is the radius of DNA. $d_s$These values yield a maximum drag force of only 0.7 fN

The hydrodynamic drag on the section of DNA threading through the portal-motor channel can also be estimated as described by Gabashvili and Grosberg: $F_{port} = v\ \xi_{port}$, where the drag coefficient is given by, $\xi_{\text{port}} = \frac{2\pi L_p\eta}{\ln\left(\frac{d_p}{d}\right)}$. Here, $L_p$ is the portal-motor channel length (at most 22 nm), and $d_p$ is the inner diameter of the channel, estimated to be ~2.5 nm (ref), based on the solved structure of the related phage P22 portal. These values yield $F_{port}$ = 0.02 fN.

Thus, all the estimated hydrodynamic drag forces are many orders of magnitude smaller than the 5 pN friction force resisting DNA exit in our measurements.

#### Supplemental References


[1] DN Fuller, et al., Measurements of single dna molecule packaging dynamics in bacteriophage $\lambda$ reveal high forces, high motor processivity, and capsid transformations. J. molecular biology 373, 1113–1122 (2007).
[2] S Chow, E Daub, H Murialdo, The overproduction of dna terminase of coliphage lambda. Gene 60, 277–289 (1987).
[3] JP Rickgauer, DN Fuller, DE Smith, Dna as a metrology standard for length and force measurements with optical tweezers. Biophys. journal 91, 4253–4257 (2006).
[4] JR Wenner, MC Williams, I Rouzina, VA Bloomfield, Salt dependence of the elasticity and overstretching transition of single dna molecules. Biophys. journal 82, 3160–3169 (2002).
[5] D delToro, DE Smith, Accurate measurement of force and displacement with optical tweezers using dna molecules as metrology standards. Appl. physics letters 104, 143701 (2014).
[6] T Odijk, Stiff chains and filaments under tension. Macromolecules 28, 7016–7018 (1995).
[7] M Ordyan, I Alam, M Mahalingam, VB Rao, DE Smith, Nucleotide-dependent dna gripping and an end-clamp mechanism regulate the bacteriophage t4 viral packaging motor. Nat. communications 9, 5434 (2018).
[8] M Doi, SF Edwards, SF Edwards, The theory of polymer dynamics. (oxford university press) Vol. 73, (1988).
[9] IS Gabashvili, AY Grosberg, Dynamics of double stranded dna reptation from bacteriophage. J. Biomol. Struct. Dyn. 9, 911–920 (1992).
[10] GC Lander, et al., Dna bending-induced phase transition of encapsidated genome in phage $\lambda$. Nucleic acids research 41, 4518–4524 (2013).
